# The SARS-CoV-2 variants associated with infections in India, B.1.617, show enhanced spike cleavage by furin

**DOI:** 10.1101/2021.05.28.446163

**Authors:** Thomas P. Peacock, Carol M. Sheppard, Jonathan C. Brown, Niluka Goonawardane, Jie Zhou, Max Whiteley, PHE Virology Consortium, Thushan I. de Silva, Wendy S. Barclay

## Abstract

The spike (S) glycoprotein of the SARS-CoV-2 virus that emerged in 2019 contained a suboptimal furin cleavage site at the S1/S2 junction with the sequence _681_P**RR**A**R**/S_686_. This cleavage site is required for efficient airway replication, transmission, and pathogenicity of the virus. The B.1.617 lineage has recently emerged in India, coinciding with substantial disease burden across the country. Early evidence suggests that B.1.617.2 (a sublineage of B.1.617) is more highly transmissible than contemporary lineages. B.1.617 and its sublineages contain a constellation of S mutations including the substitution P681R predicted to further optimise this furin cleavage site. We provide experimental evidence that virus of the B.1.617 lineage has enhanced S cleavage, that enhanced processing of an expressed B.1.617 S protein in cells is due to P681R, and that this mutation enables more efficient cleavage of a peptide mimetic of the B.1.617 S1/S2 cleavage site by recombinant furin. Together, these data demonstrate viruses in this emerging lineage have enhanced S cleavage by furin which we hypothesise could be enhancing transmissibility and pathogenicity.

## Introduction

Unlike its closest known relatives, the SARS-CoV-2 spike (S) protein contains a furin cleavage site at the S1/S2 junction that enhances SARS-CoV-2 replication in airway cells and contributes to virus pathogenicity and transmissibility (1–6). Pre-cleavage of the S protein in producer cells allows SARS-CoV-2 to enter target cells at the cell surface avoiding endosomal restriction factors (4, 7). However, the cleavage site of the early SARS-CoV-2 isolates that emerged in late 2019 are suboptimal, leaving the potential for evolution of variants with increased transmission as a result of an optimised cleavage site (4).

Towards the end of 2020 the SARS-CoV-2 pandemic entered a new phase with repeated emergence of ‘variants of concern’ lineages with altered viral properties such as transmissibility, pathogenicity, and antigenicity (8). The most widespread and best characterised of these variants is the B.1.1.7 lineage, first found in the UK, which has increased transmissibility and pathogenicity compared to other circulating strains (9–11). We and others have previously described that the S1/S2 cleavage site of B.1.1.7 S contains a P681H mutation that enhances post-translational S1/S2 cleavage during virus budding (12, 13). Other widely circulating variants that arose around the same time include the B.1.351 and P.1 lineages, first found in South Africa and Brazil, respectively, that show antigenic escape but do not contain alterations at the furin cleavage site (14). As of May 2021, an increasing number of variant lineages have been described, one of which is the B.1.617 lineage. The emergence of this lineage in India coincided with a period of record disease burden across the country, leading to partial collapse of its health infrastructure (15). Early evidence from the UK suggests one B.1.617 sublineage (B.1.617.2) likely has enhanced transmissibility, comparable to, or greater than B.1.1.7 (16). B.1.617 and its sublineages contain several S mutations, some shared with other variants and associated with antigenic escape (see Table 1). One S substitution shared by all B.1.617 sublineages is P681R which we hypothesise further optimises the furin cleavage site (_681_P**RR**A**R**/S_686_ to _681_***R*RR**A**R**/S_686_, Figure 1a). In this report we characterise the impact of P681R on the S1/S2 cleavage site.

**Table 1.** Spike Mutational profiles of B.1.617 sublineages and B.1.1.7

| Lineage | Spike mutations |
| --- | --- |
| B.1.617.1 | T95I*, G142D, E154K, L452R, E484Q, D614G, P681R, Q1071H |
| B.1.617.2 | T19R, G142D, Δ156-157/R158G, L452R, T478K, D614G, P681R, D950N |
| B.1.617.3 | T19R, Δ156-157/R158G, L452R, E484Q, D614G, P681R, D950N |
| B.1.1.7 | Δ69-70, Δ144, N501Y, A570D, D614G, P681H, T716I, S982A, D1118H |

\*Mutation found in most, but not all isolates of this sublineage

**Figure 1.**
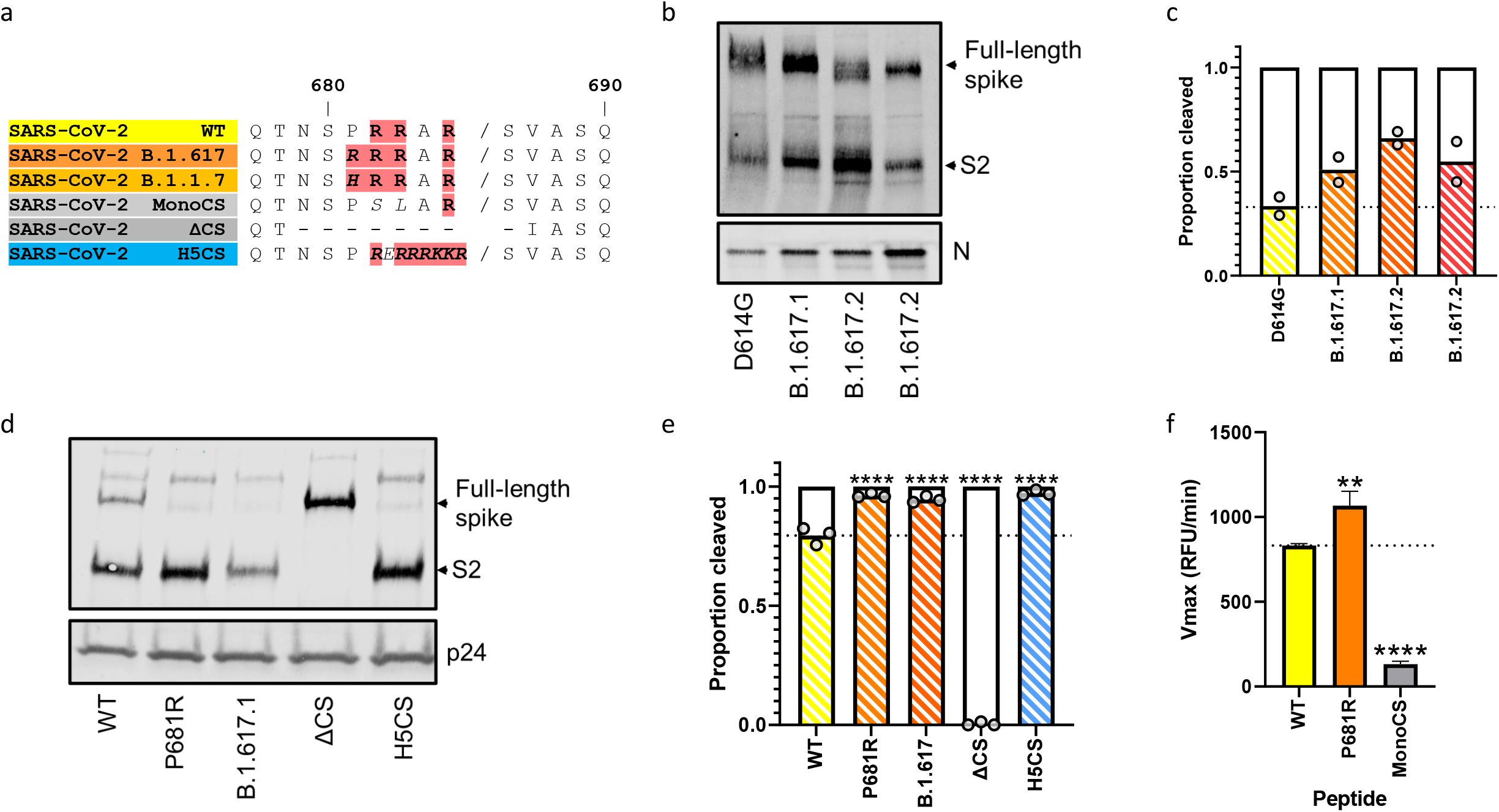
P681R results in enhanced furin cleavage of the SARS-CoV-2 B.1.617 spike protein. (a) Primary sequences of SARS-CoV-2 S1/S2 cleavage sites used throughout this study. Basic residues shown in bold and red, changes from ‘WT’ shown in italics. Numbers indicate spike residues in primary sequence (equivalent to Wuhan-Hu-1 reference sequence). (b) Western blot analysis of spike cleavage of concentrated B.1.238 (D614G) and B.1.617 (P681R containing) SARS-CoV-2 isolates. Levels of nucleocapsid (N) protein shown as loading control. (c) Densitometry analysis of the western blot from part (b). Densitometry measured using ImageJ. Points indicate two technical repeats from the same concentrated virus stocks. (d) Western blot analysis of concentrated pseudovirus containing different SARS-CoV-2 spike mutants. Levels of lentiviral p24 antigen shown as loading control. Representative blot shown of N=3 independent repeats. (e) Densitometry analysis of pseudovirus spike cleavage (from part d). Each dot indicates one completely independently produced and concentrated pseudovirus preparation (N=3). Data plotted as mean with individual repeats shown. The band corresponding to uncleaved Spike was determined by comparing the size to ΔCS which is unable to be cleaved by furin as previously described (4). Statistics performed with one-way ANOVA with multiple comparisons against the WT. \*\*\*\**P* ≤ 0.0001. (f) Cleavage of SARS-CoV-2 spike S1/S2 fluorogenic peptide mimetics by recombinant furin. Plotted as maximum enzymatic activity (Vmax). All assays performed in technical triplicate (N=3) with a representative repeat from three completely independent repeats (N=3) shown. Graph plotted as mean + Standard deviation. One-way ANOVA with multiple comparisons against the WT plotted on the graph. **0.01 ≥ *P* > 0.001; \*\*\*\**P* ≤ 0.0001.

## Results and discussion

To investigate whether the S protein of B.1.617 undergoes a higher degree of post-translational cleavage at S1/S2 than previously circulating strains, we isolated several B.1.617 lineage viruses (1 x B.1.617.1 and 2 x B.1.617.2) and compared their S1/S2 cleavage to that of a previously circulating strain of lineage B.1.238, which contains only D614G. The B.1.617 lineage S proteins were all more highly cleaved (≥50% cleaved), with a higher proportion of cleaved S2 and a lower proportion of full-length S detectable than the control virus (~33% cleaved) (Figure 1b, c).

To characterise which amino change in the B.1.617 S is responsible for its enhanced cleavage, we generated pseudovirus containing the SARS-CoV-2 full B.1.617.1 S and compared it to pseudovirus with D614G spike (WT). As we had previously observed, SARS-CoV-2 spike expressed on pseudovirus contains a larger proportion of cleaved spike (4). While WT S displayed both full length (~20%) and cleaved (~80%) S, B.1.617.1 S showed significantly enhanced cleavage (~95%), with an almost complete lack of full-length protein (Figure 1d,e). P681R alone (on a D614G backbone) was sufficient to convey this phenotype (~96% cleaved), with cleavage enhanced to a similar level as for a previously described S protein carrying the fully optimised furin cleavage site from an H5N1 avian influenza virus haemagglutinin (~97% cleaved) (4). This suggests P681R alone is responsible for the enhanced S cleavage seen in the B.1.617 lineages viruses.

We then performed assays to determine whether the optimised cleavage site found in the B.1.617 S enables better cleavage directly by furin. We measured the ability of recombinant furin to cleave fluorescently labelled peptides corresponding to the S1/S2 cleavage site of SARS-CoV-2 testing peptides containing 681P (WT), 681R, or a monobasic mutant (monoCS) whereby two of the arginines are substituted to non-basic residues (see Figure 1a) (4). As expected, monoCS was poorly cleaved by recombinant furin compared to the WT peptide which was efficiently cleaved by furin as previously described (Figure 1f)(2). P681R significantly enhanced the ability of furin to cleave the peptide confirming that the arginine substitution is responsible for the enhanced cleavage of the B.1.617 S protein.

To conclude, we speculate that enhanced S1/S2 cleavage seen in B.1.617 and B.1.1.7 (which contains P681H (12)) may be contributing to the enhanced transmissibility of these SARS-CoV-2 variants. As well as B.1.1.7 and B.1.617, several other emerging lineages contain mutations in the furin cleavage site (8). We would advise that these lineages be kept under close monitoring for any early evidence of more rapid transmission or higher pathogenesis.

## Materials and methods

### Cells and viruses

Vero E6-ACE2-TMPRSS2 (Glasgow University)(17), were maintained in DMEM, 10% FCS, 1x non-essential amino acids, 200 μg/ml hygromycin B (Gibco) and 2mg/ml G418 (Gibco). Cells were kept at 5% CO_2_, 37°C.

Upper respiratory tract swabs used to isolate viruses were collected for routine clinical diagnostic use and sequenced using the ARTIC network protocol (https://artic.network/ncov-2019) to confirm the presence of B.1.617 lineage virus, under approval by the Public Health England Research Ethics and Governance Group for the COVID-19 Genomics UK consortium (R&D NR0195). Virus was isolated by inoculating 100 μL of neat swab material onto Vero cells, incubating at 37°C for 1 h before replacing with growth media supplemented with 1x penicillin/streptomycin and 1x amphotericin B. Cells were incubated for 5-7 days until cytopathic effect was observed. Isolates were passaged a further two times in Vero E6-ACE2-TMPRSS2 cells (17), the supernatant clarified by centrifugation and concentration for western blot analysis viruses by centrifuging in an Amicon^®^ Ultra-15 Centrifugal Filter Unit followed by an Amicon^®^ Ultra-0.5 Centrifugal Filter Unit with 50 kDa exclusion size.

### Plasmids and Pseudovirus

The B.1.617.1 plasmid was generated from a previously described codon-optimised SARS-CoV-2 spike plasmid (Wuhan-hu-1)(18), using the QuikChange Lightning Multi Site-Directed Mutagenesis kit (Agilent). Pseudovirus was generated and concentrated as previously described (4). All spike expression plasmids used in this study contain D614G and K1255*STOP (that results in deletion of the C-terminal cytoplasmic tail of spike containing the endoplasmic retention signal, aka the Δ19 spike truncation).

### Western Blotting

Virus or pseudovirus concentrates were lysed in 4x Laemmli buffer (Bio-rad) with 10% β-mercaptoethanol and run on SDS-PAGE gels. After semi-dry transfer onto nitrocellulose membrane, samples were probed with mouse anti-p24 (abcam; ab9071), rabbit anti-SARS spike protein (NOVUS; NB100-56578), or rabbit anti-SARS-CoV-2 nucleocapsid (SinoBiological; 40143-R019). Near infra-red (NIR) secondary antibodies, IRDye^®^ 680RD Goat anti-mouse (abcam; ab216776) and IRDye^®^ 800CW Goat anti-rabbit (abcam; ab216773) were subsequently used to probe membranes. Western blots were visualised using an Odyssey Imaging System (LI-COR Biosciences).

### Peptide cleavage assays

The peptide cleavage assay was adapted from the protocol by Jaimes et al (2, 19). Briefly fluoregenic peptides were synthesised (Cambridge research biochemicals) with the sequences TNSPRRARSVA (WT), TNSRRRARSVA (P681R) and TNSPSLARSVA (monoCS) and, N-terminally conjugated with the fluorophore 5-Carboxyfluorescein (FAM) and the C-terminal quencher 2,4-Dinitrophenyl.

Each peptide was tested for its ability to be cleaved by recombinant furin (10 U/mL; NEB; P8077) in a buffer of 100 mM HEPES, 0.5% Triton X-100, 1mM CaCl_2_, 1 mM β-mercaptoethanol, pH 7.5. Assays were performed in triplicate at 30°C and fluorescence intensity was measured at wavelengths of 485 and 540 nm every 1 minute for 1 hour using a FLUOstar Omega plate reader (BMG Labtech). Vmax was then calculated.

## Acknowledgements and funding

The authors would like to thank Dr Matthew Turnbull and Dr Suzannah Rihn of the MRC-University of Glasgow Centre for Virus Research (CVR) for sharing their Vero E6-ACE2-TMPRSS2 cells. Thanks to members of the PHE Virology Consortium; Angie Lackenby, Shahjahan Miah, Steve Platt, Joanna Ellis, Maria Zambon and Christina Atchison, as well as PHE field staff for collection of infectious swabs and sequence information used in this work.

Sequencing of SARS-CoV-2 clinical samples was undertaken by the Sheffield COVID-19 Genomics Group as part of the COG-UK CONSORTIUM and supported by funding from the Medical Research Council (MRC) part of UK Research & Innovation (UKRI), the National Institute of Health Research (NIHR) and Genome Research Limited, operating as the Wellcome Sanger Institute.

This work was supported by the G2P-UK National Virology Consortium funded by UKRI.

